# Effect of nitrite on neural activity in the healthy brain

**DOI:** 10.1101/535336

**Authors:** Edit Franko, Martyn Ezra, Douglas C Crockett, Olivier Joly, Kyle Pattinson

## Abstract

**Background:** Nitrite is a major intravascular store for nitric oxide. The conversion of nitrite to the active nitric oxide occurs mainly under hypoxic conditions to increase blood flow where it is needed the most. The use of nitrite is, therefore, being evaluated widely to reduce the brain injury in conditions resulting in cerebral hypoxia, such as cardiac arrest, ischaemic stroke or subarachnoid haemorrhage. However, as it is still unknown how exogenous nitrite affects the brain activity of healthy individuals, it is difficult to clearly understand how it affects the ischaemic brain.

**Objective:** Here we performed a double-blind placebo-controlled crossover study to investigate the effects of nitrite on neural activity in the healthy brain.

**Methods:** Twenty-one healthy volunteers were recruited into the study. All participants received a continuous infusion of sodium nitrite (0.6mg/kg/h) on one occasion and placebo (sodium chloride) on another occasion. Electroencephalogram was recorded before the start and during the infusion. We computed the power spectrum density within the conventional frequency bands (delta, theta, alpha, beta), and the ratio of the power within the alpha and delta bands. We also measured peripheral cardiorespiratory physiology and cerebral blood flow.

**Results:** We found no significant effect of nitrite on the power spectrum density in any frequency band. Similarly, the alpha-delta power ratio did not differ between the two conditions. However, nitrite infusion decreased the mean blood pressure and increased the methaemoglobin concentration in the blood.

**Conclusion:** Our study shows that exogenous sodium nitrite does not alter the electrical activity in the healthy brain. This might be because the sodium nitrite is converted to vasoactive nitric oxide in areas of hypoxia, and in the healthy brain there is no significant amount of conversion due to lack of hypoxia. However, this lack of change in the power spectrum density in healthy people emphasises the specificity of the brain’s response to nitrite in disease.

## 1. Introduction

Nitric oxide (NO)^1^, also known as endothelium-derived relaxing factor, is an intrinsic vasodilator and signaling molecule [1]. It is synthesized from L-arginine by Nitric Oxide Synthase, an enzyme that is present in the endothelium, neurons, and macrophages [2]. NO has an important role in the regulation of the vascular tone and the maintenance of blood flow [3][4], and the adaptation of the local cerebral blood flow to the neural activity, so-called neurovascular coupling [5][6]. However, as it has a short half-life, it has to be continuously produced from its precursors [7]. Nitrite is a precursor molecule, that can be stored in the blood and transported to areas that need NO [8].

Nitrite is a major intravascular store for nitric oxide [9][10][11]. It is produced from dietary nitrate by commensal salivary bacteria in the mouth [12][13]. Nitrite can be reduced to NO by mitochondria [14], by xanthine oxidase [15], and most importantly, by deoxygenated haemoglobin [7][8], therefore producing NO in areas of low oxygen concentration. Dietary nitrate from vegetables, especially from beetroot, lettuce, and spinach [16] also has beneficial physiological effects via conversion to nitrite and then to NO. It was shown that the consumption of nitrate reduced the blood pressure [17][18][19]. It also enhanced tolerance to exercise [20][21] and improved muscle oxygenation [22]. A recent study even showed improved cognitive performance after a single bottle of beetroot juice [23]. A potential therapeutic effect of the nitrite has been shown in various animal models: intravenous sodium nitrite reduced the damage to the heart [24], liver [25] and brain after ischaemia/reperfusion [26], and prevented delayed cerebral vasospasm after subarachnoid haemorrhage in non-human primates [27].

Based on the beneficial physiological effects of nitrates and nitrites on healthy people and in animal models, studies started to evaluate the effects of nitrites in patients with different diseases. The first step was to prove the safety of such a therapy. To this end, Pluta and colleagues [28] administered intravenous sodium nitrite to healthy volunteers for 48h. They found that the maximal tolerated dose was 266.9mcg/kg/h, whereas the dose of 445.7mcg/kg/h caused an asymptomatic increase in methaemoglobin level and a transient decrease in the mean arterial blood pressure. Dezfulian and colleagues [29] performed a safety and feasibility study on patients surviving out-of-hospital cardiac arrest. In their final analysis, they found that 60mg of sodium nitrite did not have significant adverse effects [30]. Similarly, Oldfield and colleagues [31] in a phase IIa study showed that sodium nitrite can safely be administered to critically ill patients after subarachnoid haemorrhage without toxicity and systemic hypotension.

Despite the increasing use of nitrite and nitrate in both healthy people [32] and patients with various diseases [33][34], there was only one study that evaluated the effect of nitrite on the neural activity. Garry and colleagues [35] combined the infusion of sodium nitrite with electroencephalography to assess the brain’s response to the nitrite after subarachnoid haemorrhage. They found that the nitrite infusion changed the brain activity of patients who recovered well without developing further cerebral ischaemia. However, in order to understand better whether the effects are specific to the injured brain, we need to know what the physiological response is to exogenous nitrite.

Here we conducted the first study that examines the effect of nitrite on neural activity in the brain of healthy adults using quantitative electroencephalographic measurements.

## 2. Materials and Methods

### 2.1. Participants

Healthy volunteers between the ages of 18–80 years old without any history of neurological or psychiatric disease were eligible to participate in this study. Written informed consent was obtained from all of them prior the recordings. The study was approved by the South Central Oxford C NHS Health Research Authority Ethics Committee 12/SC/0366.

### 2.2. Study Design

#### 2.2.1. Electroencephalography

Each participant underwent continuous electroencephalographic (EEG)^2^ monitoring (Porti 7 system, Twente Medical Systems International, Oldenzaal, The Netherlands) on two occasions. Each recording contained around 15 minutes of baseline resting EEG with closed eyes. The baseline was followed by one hour of resting EEG with closed eyes combined with continuous intravenous infusion of either sodium nitrite or sodium chloride (placebo). Both the investigator and the participant were blinded to the content of the infusion. The two recording sessions were separated by at least two weeks. The order of sodium nitrite and placebo infusion was randomly allocated using sealed envelopes. We used a simplified electroencephalographic montage, limited by the capacity of the Porti system and the number of electrodes we could use in our patient population included in a similar study. Twelve unipolar EEG electrodes were used at the following positions defined according to the International 10–20 system: Cz, Fz, Pz, Fp1, Fp2, F3, F4, P3, P4, T3, T4, and Oz. The ground was placed close to the nasion and the references were places on the mastoids. EEG data were digitized at a sampling rate of 512 Hz (OpenViBE [36]).

#### 2.2.2. Infusion

The continuous infusion (sodium nitrite (0.6mg/kg/h) or 0.9% sodium chloride (placebo)) started after about 15 minutes of baseline EEG recording at a constant rate of 0.12ml/kg/h and continued for one hour. The dosing schedule was developed as a compromise between ensuring adequate delivery of cerebral NO and minimization of cardiovascular effects in the patient population.

#### 2.2.3. Physiologic Measurements

Participants underwent intermittent Transcranial Doppler (TCD)^3^ monitoring. Insonation of the middle cerebral artery M1 segment was performed unilaterally during the baseline recording and twice during the infusion (around 20 and 40 minutes after the onset of infusion) using color-coded duplex ultrasound (2-Mz probe; EZ-Dop GmbH, Singen, Germany). End-tidal carbon dioxide (CO_2_)^4^, end-tidal oxygen (O_2_)^5^ (Respiratory Gas Analyzer, ADinstruments Ltd, Oxford, UK), and pulse oximetry were recorded continuously and collected on a Power-1401 data acquisition interface (Cambridge Electronic Design, Cambridge, UK). As the drug might slightly reduce the blood pressure and increase the methaemoglobin level (Masimo Rainbow Pulse CO-Oximeter, Switzerland) in the blood, both of these parameters were non-invasively monitored regularly during the EEG recording (during the baseline, at 5, 10,15, 30 minutes after the start of the infusion and at the end of the infusion).

### 2.3. Data analysis

#### 2.3.1. Quantitative Electroencephalographic Analysis

Preprocessing was carried out with custom-written Python scripts using MNE software package [37]. Datasets were re-referenced to the average of the mastoid reference electrodes, then bandpass filtered between 0.5 Hz and 40 Hz using a finite impulse response function. The signal was divided into two second epochs with no overlap. Epochs, containing artefacts were removed automatically by thresholding the signal (peak-to-peak amplitude: 0.15mV). Power spectrum density was calculated between 0.5 and 25 Hz on each artefact-free epoch using multitapers with a resolution of 0.5 Hz. The power spectrum density was then averaged across channels, within the first and second 30 minutes of the infusion and within conventional frequency bands: delta: 1-3.5Hz, theta: 4-7.5Hz, alpha: 8-12Hz, beta: 12.5-25Hz. We also computed the alpha-delta ratio by dividing the power in the alpha band with the power in the delta band and taking the square root of it [35]. For statistical analysis, we used both the raw power and the percentage of power change (%change) of the normalized EEG. The %change for each frequency was calculated as:

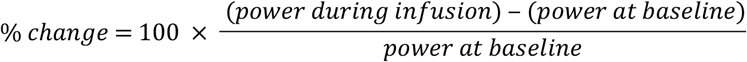

The raw EEG power and %change were averaged across subjects and compared between placebo and sodium nitrite infusion.

#### 2.3.2. Physiologic Data Analysis

We used custom-written Python codes for the analyses of the physiological measurements. End-tidal O_2_ and CO_2_ were recorded and digitalized at a sampling frequency of 100Hz. For the O_2_, the end-tidal value of each breath was identified as the local minimum within 100 neighbouring values. To exclude potential artefacts, only values below 19 kPa were then kept for further analysis. These end-tidal O_2_ values were averaged within the baseline period and 30 minute blocks during the infusion period similar to the EEG analysis. The end-tidal CO_2_ value was identified as the local maximum within 100 neighbouring values. To exclude potential artefacts, only values above 1.7 kPa were kept for further analysis. Similarly to the end-tidal O_2_, the end-tidal CO_2_ values were then averaged within the baseline and the first and the second 30 minutes of infusion.

Transcranial Doppler measurement: artefact-free parts from the 3 blocks of TCD recordings (one during the baseline, two during the infusion periods) were selected for further analysis. The systolic velocity (SV)^6^ within each heart cycle was defined as the local maximum within 50 neighbouring values. The diastolic velocity (DV)^7^ was defined as the local minimum within 50 neighbouring values. From these two values, the mean velocity (MV)^8^, the pulsatility index (PI)^9^ and the resistance index (RI)^10^ were calculated within each heart cycle using the following formulas:

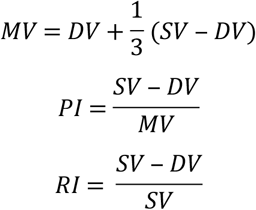

These values were then averaged within the three recording time blocks (baseline, and two infusion blocks).

#### 2.3.3. Statistical Analysis

We analysed both the raw power and the %change after normalizing to the baseline as these different ways to account for how the baseline power might influence the results. To test our hypothesis whether sodium nitrite infusion changes the raw power of the brain activity, we used a linear model. The dependent variable was the power during the infusion, the independent variable was the type of infusion (nitrite or placebo). We included the power during the baseline and the age as potential confounders into the model. As normalization is another way to account for the variability during the baseline, we also compare the %change during nitrite infusion with %change during placebo infusion using a paired t-test. Both the linear model and the paired t-test were repeated separately for each frequency band (delta, theta, alpha, beta) and for the alpha-delta ratio.

The end-tidal O_2_ and CO_2_ values were normalized by subtracting the baseline values from the values during the infusion. A paired t-test was then used to compare the values between the nitrite and the placebo infusion separately for the first and the second time block.

The five TCD measurements, SV, DV, MV, PI, and RI, were normalized by subtracting from the infusion period the corresponding value in the baseline. A paired t-test was then used to look for a significant difference between the nitrite and the placebo infusion.

Blood pressure and methaemoglobin values were normalized by subtracting the baseline value separately from each measurement during the infusion period. The normalized values were then compared between the nitrite and placebo infusion using paired t-tests within each time point (5, 10, 15, 30 minutes after the start of the infusion and at the end of it). The systolic, diastolic and mean blood pressure values were analysed separately.

For each statistical test, the alpha level was set at 0.05 after correcting for multiple comparisons (Bonferroni correction).

## 3. Results

### 3.1. Participants

Twenty-one healthy people were included in the study. The mean age was 46.4 ± 15.6 (SD) years (range: 22-68), seven of them were females. Two participants attended only one visit, therefore there were 40 EEG recordings (19 sessions with sodium nitrite infusion, 21 sessions with placebo infusion).

### 3.2. EEG

The result of the time-frequency analysis of one subject is shown in Figure 1 for both the placebo (Figure 1A) and the nitrite (Figure 1B) infusion. We used normalization with the baseline to remove the effect of baseline difference in the display. When comparing the effect of nitrite to placebo infusion on the EEG power using baseline power as a predictor variable for all the subjects, the linear model did not show any significant difference within any of the four frequency bands, neither in the alpha-delta ratio. Supplementary Figure 1 shows the power in the four frequency bands and in the alpha-delta ratio in the baseline during the first and second 30 minutes of infusion. However, we found a significant effect of baseline on the power in each frequency band. Supplementary Table 1 shows the t and p-values for the different frequency bands. There was a weak effect of age on the power in the delta band during the first half of the infusion; however, it did not reach a significant level after correction for multiple comparisons.

**Figure 1.**
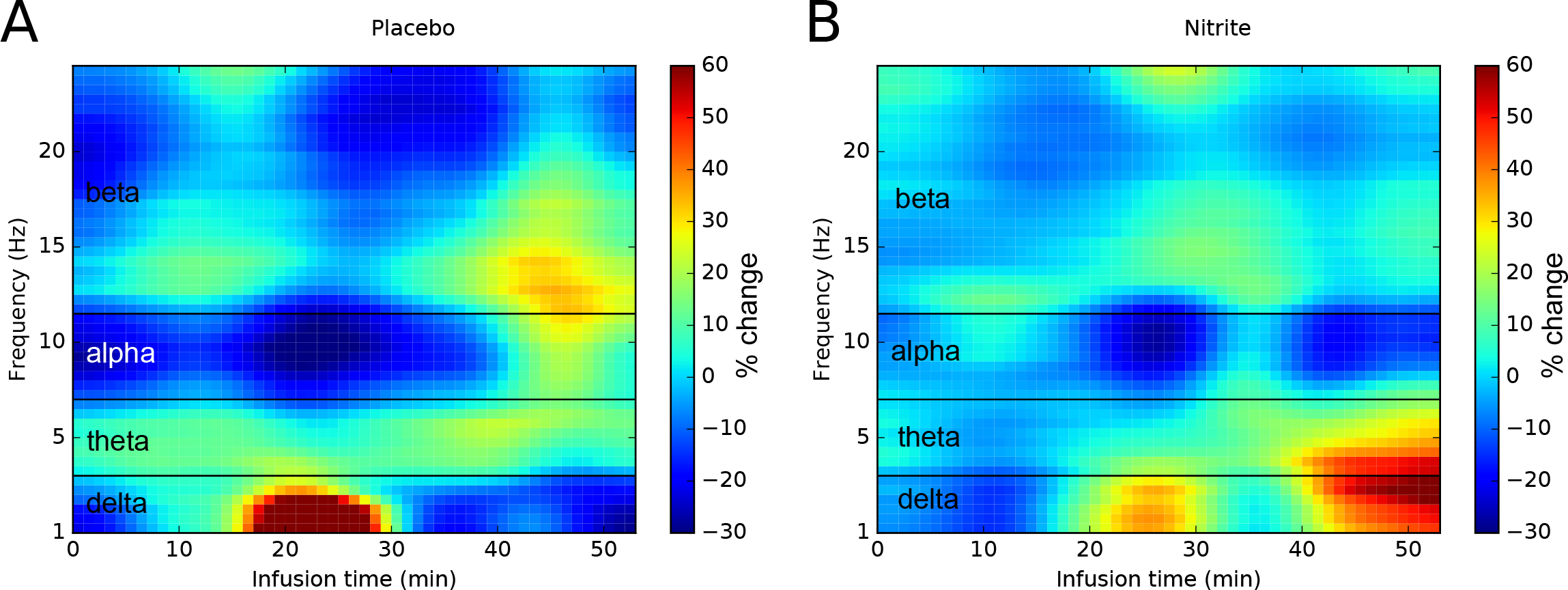
The power spectrum density of one participant after normalisation with the baseline power. A, placebo infusion. B, nitrite infusion. The power spectrum density was averaged within 1-minute blocks of the infusion period and smoothed for visualisation (Gaussian).

When we looked at the data after normalization with the baseline power, we found similar results. Using a paired t-test to compare the % change in power between the nitrite and placebo infusion, we could not find any significant difference. The results of the t-tests are shown in Table1. There was a trend in the beta band (mean difference: 8.66, p=0.07 uncorrected) indicating a slightly increased beta power during the nitrite infusion. Figure 2 shows the % of power change in the different frequency bands for the nitrite and placebo infusions for the first and second 30 minutes separately. Even towards the end of the infusion, the power was similar for both types of infusions.

**Table 1.**
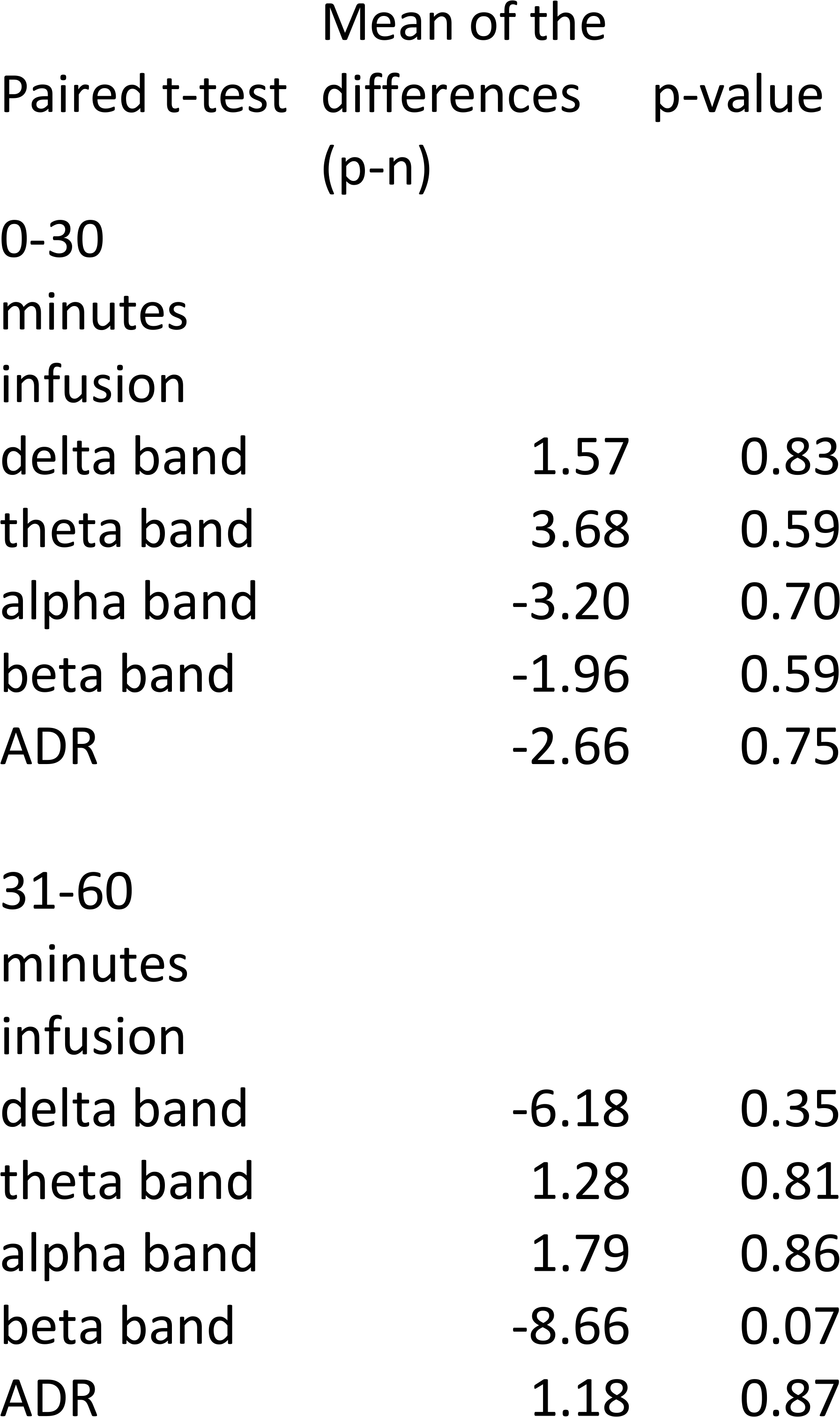
Results of the paired t-test (mean of the differences between placebo and nitrite, and p-values) on the power change in the different frequency bands within the first and the second 30 minutes of infusion.

**Figure 2.**
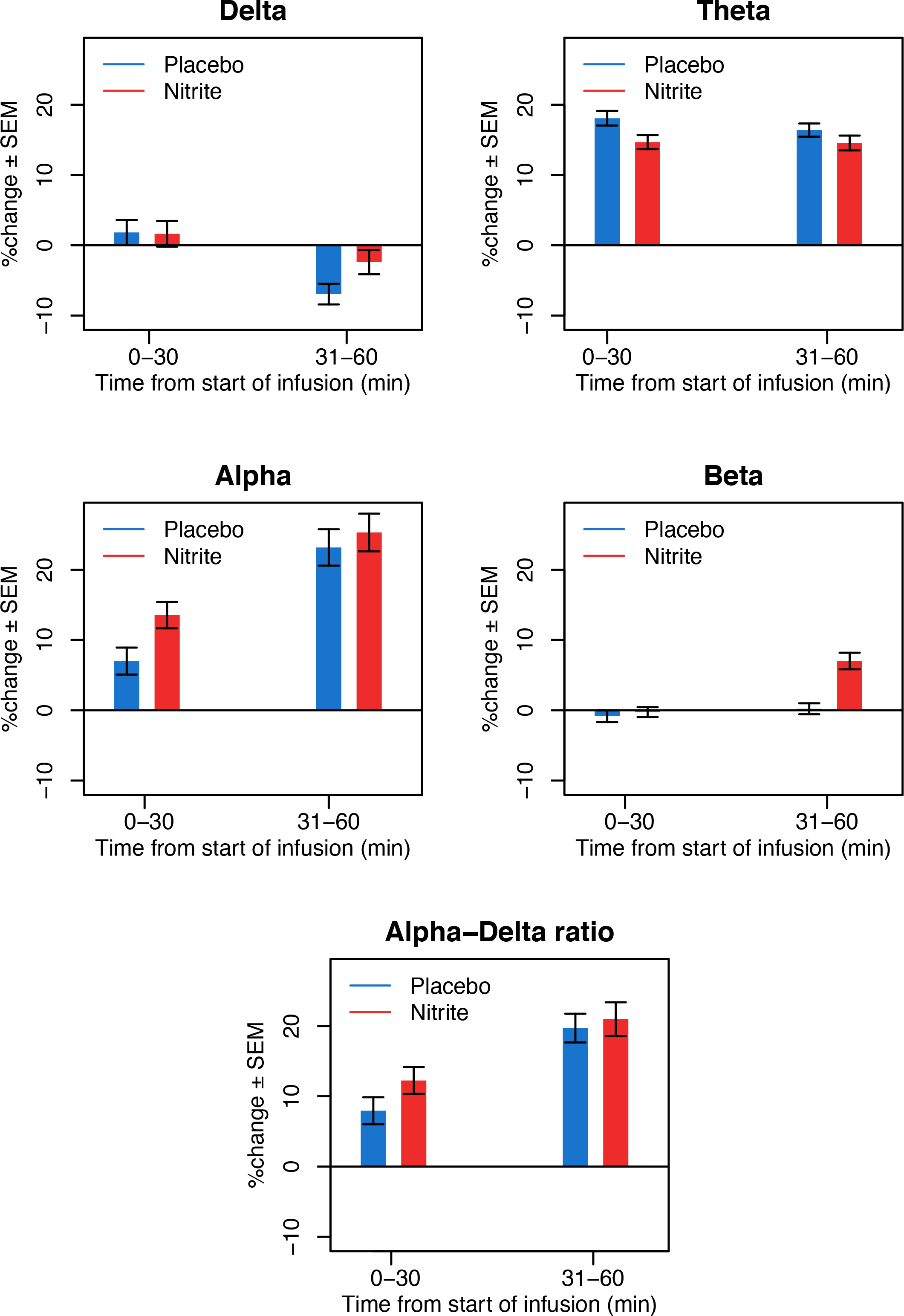
The normalised power spectrum density in the separate frequency bands (delta, theta, alpha, beta) and in the alpha/delta ratio. The power was averaged within the first and the second 30 minutes of infusion. SEM: standard error of the mean.

### 3.3. Physiological measurements

In one session, the end-tidal gases were not recorded due to technical issues, therefore there were 19 data during nitrite and 20 data during placebo infusion. There were six sessions when we could not get a good quality TCD signal. Therefore, together with the two missed sessions, there were eight missing TCD data leaving us with 14 data during nitrite infusion and 20 data during placebo infusion.

There was no significant difference in the end-tidal O_2_ and CO_2_; and any of the TCD measurements (SV, DV, MV, PI, and RI) between the nitrite and placebo infusions at any time points. The mean of the differences (placebo-nitrite) and the p-values for the last time points are shown in Table 2 as we expected stronger difference towards the end of the infusion time due to the accumulation of the nitrite in the blood. Supplementary figures show the mean values in the first and second 30 minutes of infusion (O_2_ and CO_2_: Supplementary Figure 2, TCD: Supplementary Figure 3).

**Table 2.** Results of the paired t-test (mean of the differences between placebo and nitrite, and p-values) on the transcranial Doppler measurements and on the end-tidal oxygen and carbon dioxide values within the first and the second 30 minutes of infusion.

| Paired t-test | Mean of the<br>differences<br>(p-n) | p-value |
| --- | --- | --- |
| 0-30 minutes<br>infusion |  |  |
| End-tidal O2 | -0.07 | 0.53 |
| End-tidal CO2 | 0.002 | 0.96 |
| TCD SV | 0.14 | 0.12 |
| TCD DV | 0.06 | 0.14 |
| TCD MV | 0.09 | 0.12 |
| TCD PI | -0.02 | 0.67 |
| TCD RI | -0.006 | 0.73 |
| 31-60 minutes<br>infusion |  |  |
| End-tidal O2 | 0.01 | 0.94 |
| End-tidal CO2 | -0.004 | 0.96 |
| TCD SV | 0.05 | 0.40 |
| TCD DV | 0.03 | 0.48 |
| TCD MV | 0.04 | 0.44 |
| TCD PI | -0.02 | 0.80 |
| TCD RI | -0.005 | 0.82 |

By the end of the session, the nitrite infusion significantly decreased all three values of the normalized blood pressure measurements, the systolic (Mean ± standard deviation: −7.4±12 mmHg), the diastolic (−6.7±8 mmHg) and the mean blood pressure (−6.9±9 mmHg). Table 3 shows the mean of the differences between placebo and nitrite infusion and the corresponding p-values for the pair-wise comparisons. There was a slight increase in systolic blood pressure five minutes after the start of the nitrite infusion (6.8 mmHg), however, the diastolic and mean blood pressure did not change. Moreover, this difference was not present at 10, 15 and 30 minutes. Figure 3A shows how the mean blood pressure changed during the time of the infusion. The systolic and diastolic blood pressure changes are shown in Supplementary Figure 4.

**Table 3.** Results of the paired t-test (mean of the differences between placebo and nitrite, and p-values) on blood pressure (BP) and methaemoglobin values. Measurements were taken 5, 10, 15, 30 and 60 minutes after the start of the infusion. Significant values are shown in bold.

|  | Mean of the<br>differences<br>(p-n) | p-value |
| --- | --- | --- |
| 5 minutes |  |  |
| Mean BP | -3.019 | 0.011 |
| Systolic BP | -6.84 | 0.03 |
| Diastolic BP | -1.36 | 0.53 |
| Methaemoglobin | 0.02 | 0.73 |
| 10 minutes |  |  |
| Mean BP | -2.05 | 0.3 |
| Systolic BP | -5.52 | 0.08 |
| Diastolic BP | -0.31 | 0.87 |
| Methaemoglobin | -0.12 | 0.23 |
| 15 minutes |  |  |
| Mean BP | -1.26 | 0.51 |
| Systolic BP | -2.94 | 0.24 |
| Diastolic BP | -0.42 | 0.84 |
| Methaemoglobin | -0.16 | 0.06 |
| 30 minutes |  |  |
| Mean BP | 3.63 | 0.08 |
| Systolic BP | 3.21 | 0.24 |
| Diastolic BP | 3.84 | 0.07 |
| Methaemoglobin | -0.44 | **0.0001** |
| 60 minutes |  |  |
| Mean BP | 6.98 | **0.003** |
| Systolic BP | 7.47 | **0.014** |
| Diastolic BP | 6.73 | **0.003** |
| Methaemoglobin | -0.82 | **8.09E-09** |

**Figure 3.**
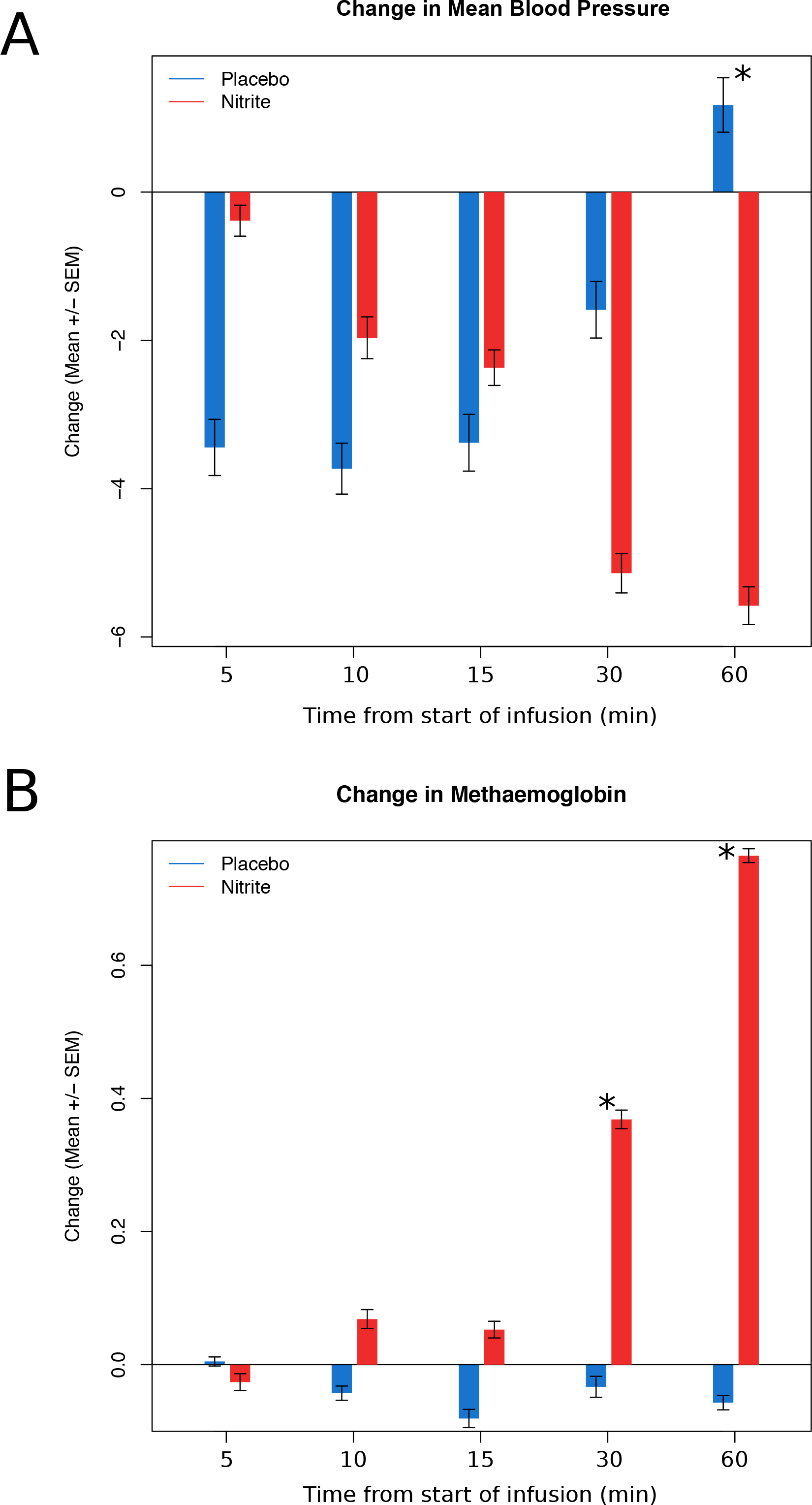
The changes over time in the mean blood pressure (A) and the methaemoglobin concentration (B). Measurements were taken 5, 10, 15, 30 and 60 minutes after the start of the infusion. The values were normalised by subtracting the baseline values. The * indicates a significant difference between placebo and nitrite infusion (p<0.05). SEM: standard error of the mean.

There was a significant increase in the methaemoglobin concentration in the blood after 30 minutes of nitrite infusion (0.44%). This increased reached about 0.82% by the end of the one-hour infusion (mean ± standard deviation methaemoglobin concentration during nitrite infusion: 1.85±0.30; during placebo infusion: 1.02±0.20). Figure 3B shows the change over time, and Table 3 shows the mean of the differences and the corresponding p-values for each time points.

## 4. Discussion

In the current study, we examined the effects of sodium nitrite infusion on the EEG of healthy adults. When analysing the power within the delta, theta, alpha and beta bands, and the ratio of the alpha and delta power, there were no significant differences between the nitrite and placebo infusions. Similarly, sodium nitrite infusion did not alter the end-tidal O_2_ and CO_2_ values and the flow in the middle cerebral artery measured by TCD. However, we did see an increase in the blood methaemoglobin concentration and a decrease in the blood pressure when administering sodium nitrite.

The change in the blood pressure and the concentration of methaemoglobin during the nitrite infusion were expected as a previous study reported asymptomatic change with 445.7mcg/kg/h nitrite infusion [28]. Even though our infusion rate was 600mcg/kg/h nitrite, which is slightly higher, the changes remained asymptomatic.

So far there have been few studies that examined the effects of exogenous nitrite and nitrate on the brain. Presley and colleagues [38] compared low versus high nitrate diet in older adults. Nitrate is converted to nitrite in the body by oral bacteria, therefore, having the same effects as direct intravenous nitrite [13]. Presley and colleagues [38] found that the high nitrate diet did not alter global cerebral perfusion, even though it increased regional cerebral perfusion in the frontal lobe white matter. The lack of change in cerebral perfusion could explain why we did not see any change in the overall neural activity. However, it does not exclude the possibility of any regional change.

Nitrite is converted to NO preferentially under hypoxic conditions to increase blood flow where it is needed the most [10]. The lack of change in the brain activity might, therefore, be due to the lack of hypoxia in healthy people’s brain. Our study population contained both young and older participants. It could be that hypoxia affects only the older population and averaging their results with the results from young people hides the effect of nitrite in this group. We, therefore, included age as a potential confounder in the linear model when comparing the effect of nitrite to placebo infusion. We found a mild effect of age on the power in the delta band in the first 30 minutes of infusion. However, it was not significant after correcting for multiple comparisons and even this mild effect was not present in the second half of the infusion. This indicates that the effect of nitrite on the EEG activity does not depend on the age.

The lack of effect might also be due to a low dose of nitrite infusion. Piknova and colleagues [39] found that the increase in the cerebral blood flow in rats was dose dependent. Especially in healthy people, where there is no hypoxia, which would increase the conversion of nitrite to NO, higher doses of nitrite might be needed to detect an increase in blood flow and neural activity.

The lack of change in brain activity could also be due to the fact that we recorded resting EEG. Nitrite is a store of NO, and it would be converted to the active molecule when there is a need for increased blood flow. As our protocol did not contain any task, there was no need for an increase in the cerebral blood flow. Previous studies showing a change in cerebral blood flow used stimulation. Aamand and colleagues [40] presented visual stimuli while measuring the blood oxygenation level dependent (BOLD)^11^ response, and found that both the latency and the amplitude of the hemodynamic response reduced after nitrate intake. Wightman and colleagues [23] activated the prefrontal cortex with cognitive tasks when they found an initial increase in the cerebral blood flow, which was followed by a reduction during the least demanding cognitive task. Only Presley and colleagues [38] found an increase in the resting blood flow in the frontal lobe white matter after nitrate intake, but they did not detect any global change. Using a task in our study might have helped reveal enhanced neurovascular coupling which could have resulted in a change in the EEG activity.

It is interesting to see that our results resemble the findings of Garry and colleagues [35] in patients at risk of deterioration. However, it helps explain the nitrite effect on patients with good outcome. These patients responded with an increase in the ratio of the alpha and delta power to nitrite infusion. It was shown previously that the reduction in blood flow could be detected by EEG as a slowing of the main background frequency [41]. Another study found that it was the delta activity that showed the strongest correlation with the regional cerebral blood flow [42]. This would suggest that patients, who showed increased alpha/delta ratio after the nitrite infusion, had increased cerebral blood flow. The lack of any change in the alpha/delta ratio in patients at risk of deterioration would then indicate no vasodilation and therefore no change in blood flow. This could be explained by severe damage to the endothelial cells that do not react anymore to the NO. This endothelial dysfunction [43] could then increase the chance to develop delayed cerebral ischaemia. However, we should not forget that NO has many different effects besides causing vasodilation in the ischaemic brain [44]. It reduces platelet aggregation and leukocyte adhesion to the vessel wall, therefore preventing microvascular plugging. NO also has a direct effect on neurons; it inhibits calcium influx into the neurons, therefore limiting glutamate neurotoxicity. It can even reduce the damage in ischaemia by scavenging reactive oxygen species. All these neuroprotective mechanisms could participate in the improvement of neural activity that was shown in patients with subarachnoid haemorrhage [35]. As the healthy brain does not have ischaemic regions, these protective mechanisms are not activated even when the nitrite concentration is increased by intravenous infusion.

## 5. Conclusion

We demonstrated in this placebo-controlled double-blind cross-over study, that sodium nitrite infusion does not affect the global brain activity in healthy adults when measured with resting quantitative EEG. Our findings serve as a base to explain the effects of nitrite in brain ischaemia, such as after subarachnoid haemorrhage.

---

**Supplementary Figure 1.**
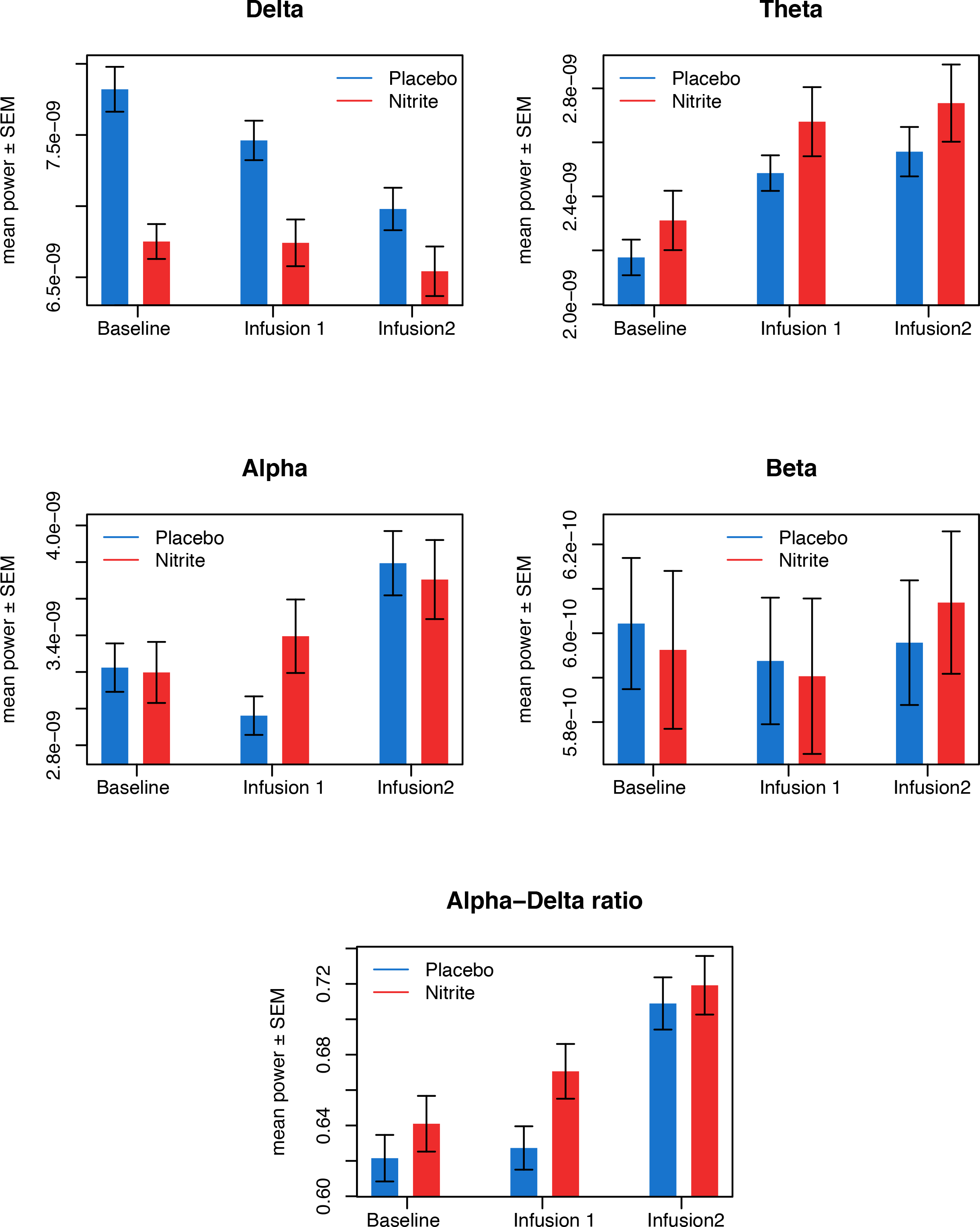
The raw power spectrum density in the separate frequency bands (delta, theta, alpha, beta) and in the alpha/delta ratio. The power was averaged within the baseline, the first and the second 30 minutes of infusion. SEM: standard error of the mean.

**Supplementary Figure 2.**
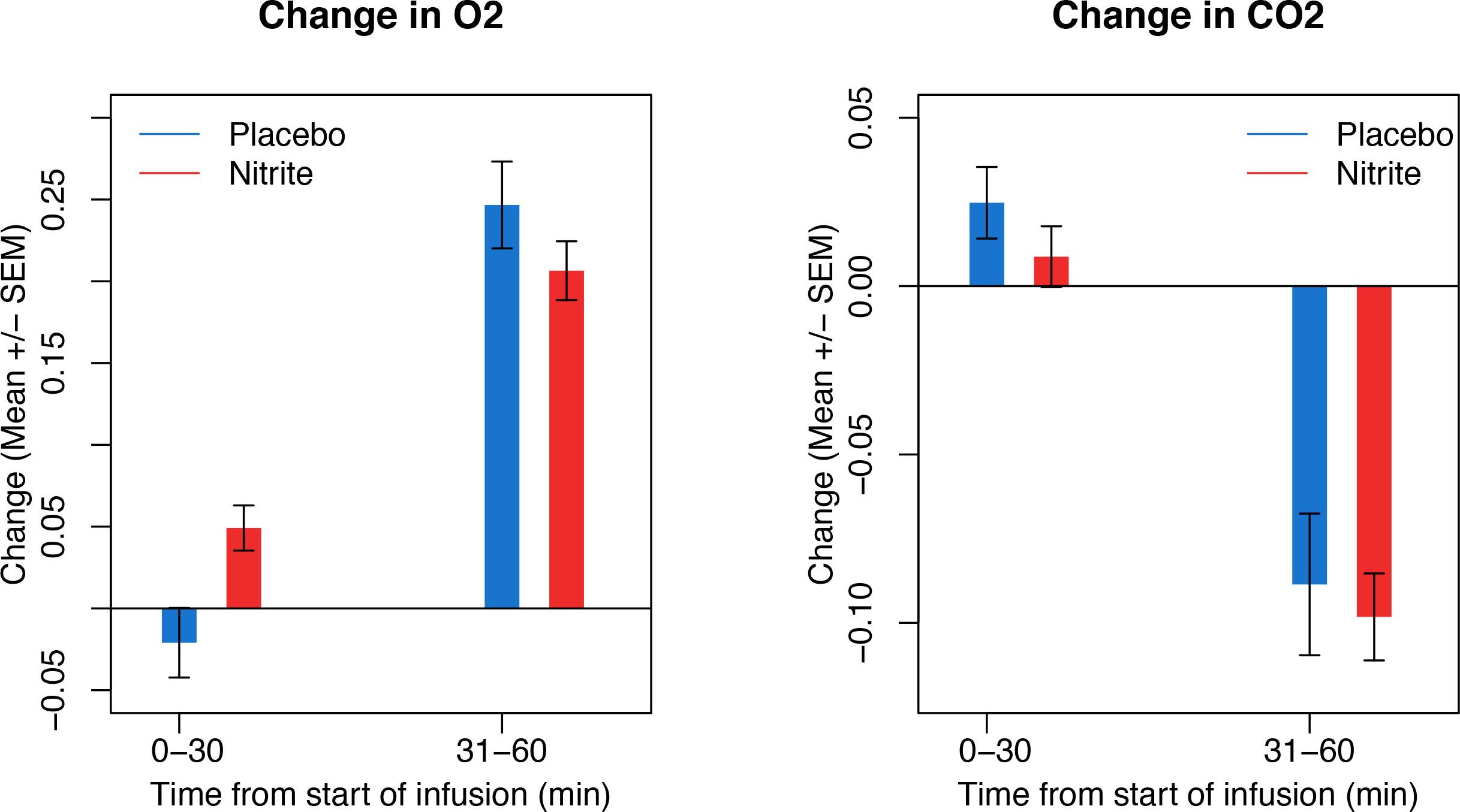
The changes over time in the end-tidal oxygen and carbon-dioxide values averaged within the first and second 30 minutes of infusion. The values were normalised by subtracting the baseline values. SEM: standard error of the mean.

**Supplementary Figure 3.**
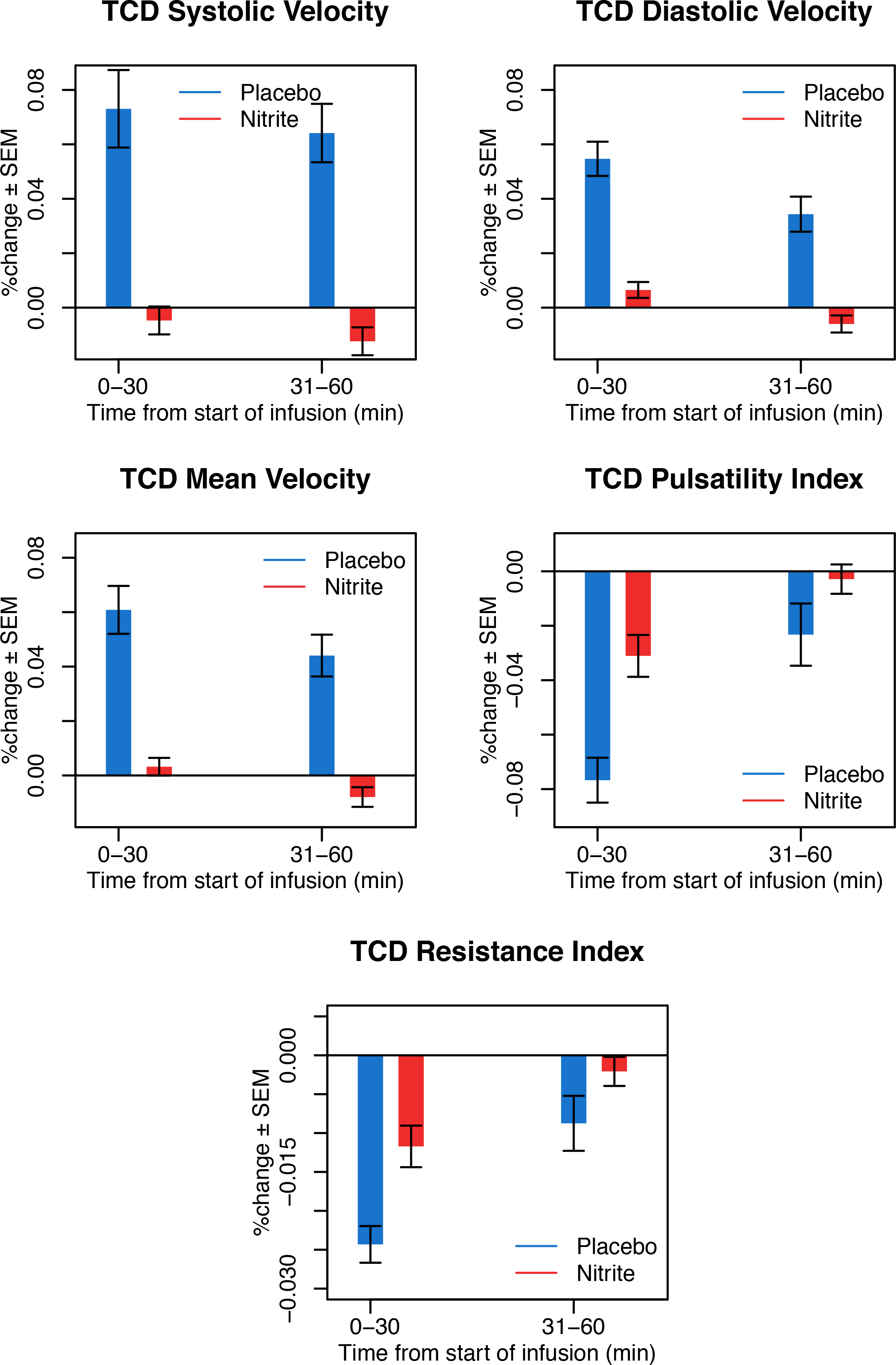
The changes over time in the transcranial Doppler measurements (systolic velocity, diastolic velocity, mean velocity, pulsatility index, resistance index) averaged within the first and second 30 minutes of infusion. The values were normalised by subtracting the baseline values. SEM: standard error of the mean.

**Supplementary Figure 4.**
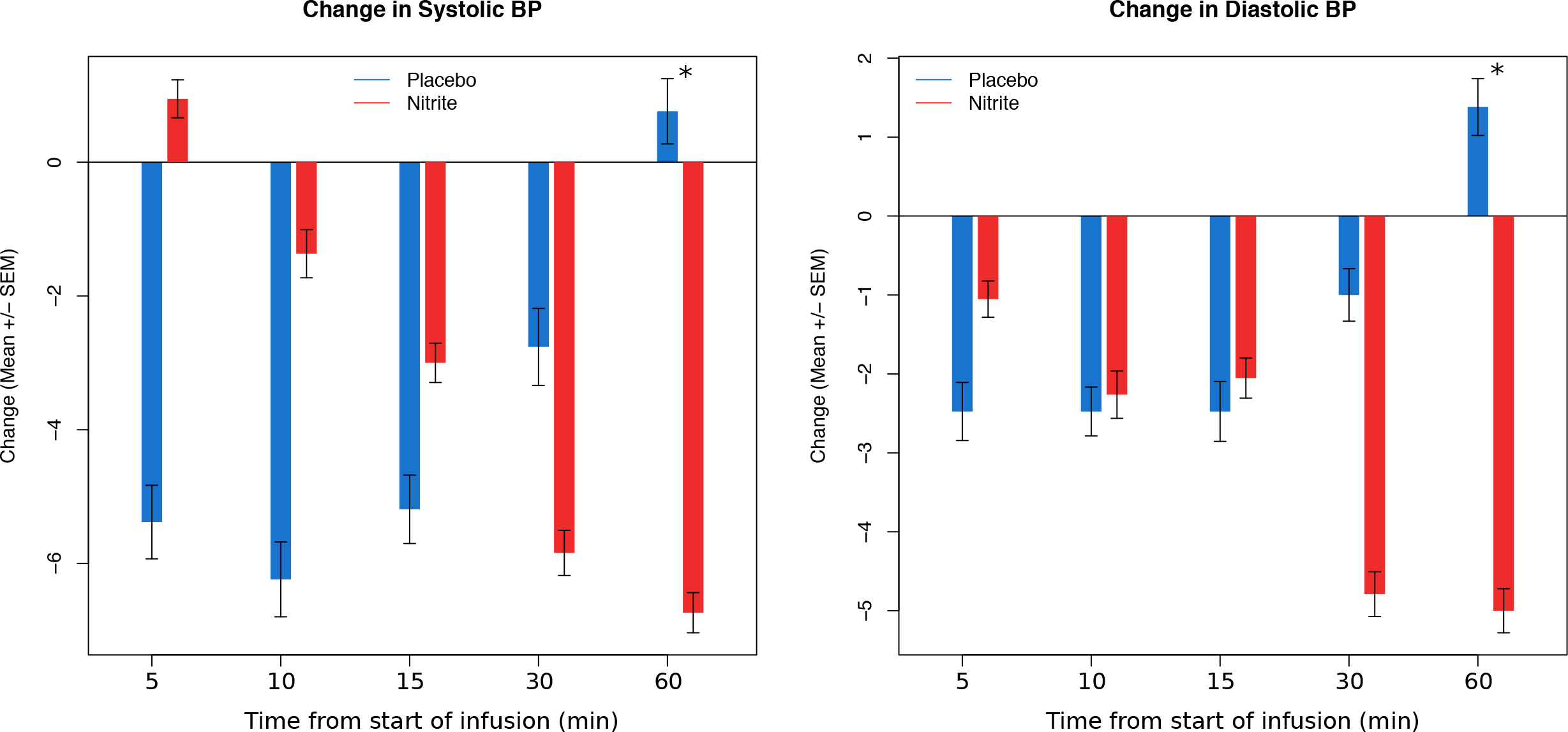
The changes over time in the systolic and diastolic blood pressure. Measurements were taken 5, 10, 15, 30 and 60 minutes after the start of the infusion. The values were normalised by subtracting the baseline values. The * indicates a significant difference between placebo and nitrite infusion (p<0.05). SEM: standard error of the mean.

**Supplementary Table 1.** Results of the linear model on the power spectrum density in the different frequency bands within the first and the second 30 minutes of infusion. Independent variables were the type of infusion (placebo or nitrite), the baseline power within the given frequency and the age. Significant values are shown in bold (p<0.005 corrected).

| Linear model | 0-30 minutes<br>infusion | t-value | p-value | 31-60 minutes<br>infusion | t-value | p-value |
| --- | --- | --- | --- | --- | --- | --- |
| Delta | type of infusion | 0.24 | 0.81 |  | -0.42 | 0.67 |
|  | baseline power | 3.05 | **0.004** |  | 3.36 | **0.001** |
|  | age | -2.72 | 0.01 |  | -1.31 | 0.19 |
| Theta | type of infusion | -0.41 | 0.68 |  | 0.01 | 0.98 |
|  | baseline power | 24.84 | **2.00E-16** |  | 19 | **2.00E-16** |
|  | age | -0.5 | 0.61 |  | 1.2 | 0.23 |
| Alpha | type of infusion | -0.45 | 0.65 |  | 0.44 | 0.65 |
|  | baseline power | 8.15 | **1.63E-09** |  | 6.28 | **3.68E-07** |
|  | age | 1.45 | 0.15 |  | 0.58 | 0.56 |
| Beta | type of infusion | -0.29 | 0.77 |  | -0.73 | 0.46 |
|  | baseline power | 14.87 | **2.00E-16** |  | 13.8 | **1.68E-15** |
|  | age | 0.2 | 0.84 |  | 1.53 | 0.13 |
| ADR | type of infusion | -0.38 | 0.7 |  | 0.27 | 0.78 |
|  | baseline power | 5.48 | **4.02E-06** |  | 4.99 | **1.74E-05** |
|  | age | 1.46 | 0.15 |  | 0.53 | 0.59 |

## Footnotes

1 *NO*, nitric oxide

2 *EEG*, electroencephalogram;

3 *TCD*, transcranial Doppler;

4 *CO*_*2*_, carbon dioxide

5 *O*_*2*_, oxygen

6 *SV*, TCD systolic velocity;

7 *DV*, TCD diastolic velocity;

8 *MV*, TCD mean velocity;

9 *PI*, TCD pulsatility index;

10 *RI*, TCD resistance index;

11 *BOLD*, blood oxygenation level dependent;

